# Zebrafish otic vesicle and mouse epididymis as model systems for studying columnar epithelial cell division

**DOI:** 10.1101/2025.10.28.683518

**Authors:** Yu Xia, Björn Perder, Alvin Gea Chen Yao, Maiko Matsui, Miaoyan Qiu, Geoffrey S. Pitt, Jingli Cao

**Affiliations:** Cardiovascular Research Institute, Weill Cornell Medicine, 1300 York Avenue, New York, NY, USA; Department of Cell and Developmental Biology, Weill Cornell Medicine, 1300 York Avenue, New York, NY, USA

**Keywords:** epithelial cell division, interkinetic nuclear migration, zebrafish otic vesicle, mouse epididymis, live imaging

## Abstract

Epithelial cell division maintains tissue architecture through coordinated nuclear migration, cell shape changes, and spindle orientation. In columnar epithelia, interkinetic nuclear migration (INM) involves apical nuclear translocation in G2 phase and basal return post-mitosis, yet its regulation remains incompletely understood in vertebrates, in part due to limited *in vivo* live-imaging models. In this methodological study, we adapted the zebrafish embryonic otic vesicle as an *in vivo* model to investigate cell division dynamics in simple columnar epithelium using high-resolution live imaging and genetic tools. We demonstrated that apical INM initiates in mid-to-late G2 and is driven by dynein, not myosin II. Mitotic rounding is achieved via actomyosin-mediated basolateral constriction while maintaining basal attachment. Inhibiting myosin II impairs rounding and planar division, causing apical retention of daughter cells, suggesting planar division ensures proper integration. We additionally analyzed the mouse epididymal epithelium, a simple columnar epithelial tissue, to allow cross-species comparison of nuclear migration dynamics. Together, these optimized *in vivo* vertebrate models uncover conserved and tissue-specific mechanisms underlying epithelial organization and function, and importantly, provide tools for deeper mechanistic dissection in the future.

## INTRODUCTION

Epithelial cell division is a highly regulated process that preserves tissue architecture and function by coordinating cell shape changes, polarity, and spindle orientation. A defining feature of epithelial tissues is their apical-basal polarity, which supports barrier function, directional transport, and tissue homeostasis^1^. The apical membrane lines the lumen, the basal membrane anchors cells to the basal lamina, and lateral membranes couple neighboring cells through tight junctions and adherens junctions. During mitosis, epithelial cells undergo extensive morphological remodeling yet maintain tissue integrity through dynamic rearrangements of cortical actin and intercellular junctions. Correct spindle orientation is critical for reintegrating daughter cells into the epithelial sheet, and its disruption can lead to developmental abnormalities and disease^2, 3^.

Columnar epithelium can be broadly classified into simple and pseudostratified columnar epithelium. In simple columnar epithelia, such as that in the stomach, intestine, airway epithelia, otic vesicle, epididymis, and prostate, cells are column-shaped with nuclei positioned near the basal lamina^4–9^. In contrast, pseudostratified epithelia, such as neuroepithelium, exhibits a “pearl-on-a-string” architecture in which cells extend thin apical and basal processes, and nuclei appear multilayered^10–13^. A hallmark of both epithelial types is interkinetic nuclear migration (INM), during which nuclei move apically during G2 and return basally following mitosis^8, 10, 11, 14–16^. Cell division in these tissues also occurs with defined geometric constraints, as mitotic spindles are typically oriented parallel to the apical surface, ensuring planar division^8, 10, 11, 17, 18^.

Mechanistic studies of INM have focused primarily on stem or progenitor cells such as within the neuroepithelium of the developing retina and neural tube. Both microtubules (MTs) and filamentous actin (F-actin) have been implicated in driving nuclear movement^10, 11, 19^, yet the relative contribution of each cytoskeletal system remains debated, likely reflecting differences in model systems or perturbation strategies. In rodent neuroepithelium, cytoplasmic dynein, a MT-bound minus-end motor, drives basal-to-apical nuclear movement, aided by apically tethered centrosomes and nuclear-anchoring proteins^10, 20–24^. Kinesin-3, a MT-bound plus-end motor, drives apical-to-basal movement, with myosin II reported to be dispensable^20^, although other studies propose a key role for myosin II in basal transit^25^. In contrast, INM in the zebrafish retina relies predominantly on myosin II in both directions^26^, and in Drosophila brain neuroepithelium, actomyosin is the primary driver of apical migration^12^. These findings highlight that although INM is driven by common cytoskeletal machinery, its regulation varies substantially across tissues and species, prompting the question of how epithelial architecture influences nuclear dynamics and division behavior. In pseudostratified epithelia such as neuroepithelium, nuclei migrate through narrow, elongated processes^10, 11, 19, 27^, whereas in simple columnar epithelia, nuclear movement during G2 may be less physically constrained. Nevertheless, mitotic rounding, during which cells transform into a spherical shape, still requires cytoplasmic displacement toward the apical surface. Thus, while INM and mitosis are shared features across columnar epithelia, mechanical constraints imposed by epithelial organization likely shape the forces and pathways that control them.

Studies of fixed *in vivo* tissues, along with live imaging in mammalian intestinal organoids, chicken neural tube slices, and *Drosophila* brain epithelia, have revealed principles of nuclear migration and mitotic behavior^12, 28–30^. However, significant gaps remain particularly in understanding how cells integrate biochemical and mechanical cues to position the nucleus, orient the spindle, determine division plane, and preserve epithelial architecture during proliferation. Commonly used *in vitro* systems often lack physiological organization and mechanical context, underscoring the need for *in vivo* vertebrate models that capture the full complexity of columnar epithelial division.

To address these gaps, we built upon work from the Megason group and others^31–33^ and adapted the zebrafish embryonic otic vesicle as an optimized *in vivo* system for studying simple columnar epithelial cell division dynamics, an application has not been explored in this context, to address the questions outlined above. The otic vesicle epithelium provides excellent optical access, high cellular resolution, and genetic manipulability, enabling deep visualization and perturbation of division events within an intact columnar tissue. We further validated key observations in the mouse epididymis epithelium, a comparable simple columnar tissue not previously examined in this context, revealing conserved features of nuclear migration and spindle orientation across vertebrates.

## RESULTS

### INM in zebrafish otic vesicle epithelium initiates in late G2

To examine nuclear behavior during cell division in the zebrafish otic vesicle epithelium, we labeled nuclei with H2B-GFP and cell membranes with RFP-F^34, 35^ by microinjecting *in vitro*-transcribed mRNAs into fertilized eggs. We focused on the ventral side of otic vesicle, where epithelial cells retain a tall, columnar morphology between 22-28 hpf (Fig. 1A, B). As expected, most nuclei remained basally positioned at steady state, but occasional apical nuclear translocations followed by mitosis were observed (Fig. 1C; Supplementary Videos S1, S2). The complete migration cycle (basal to apical and back to basal) lasted 70.4 ± 7.7 min.

**Figure 1.**
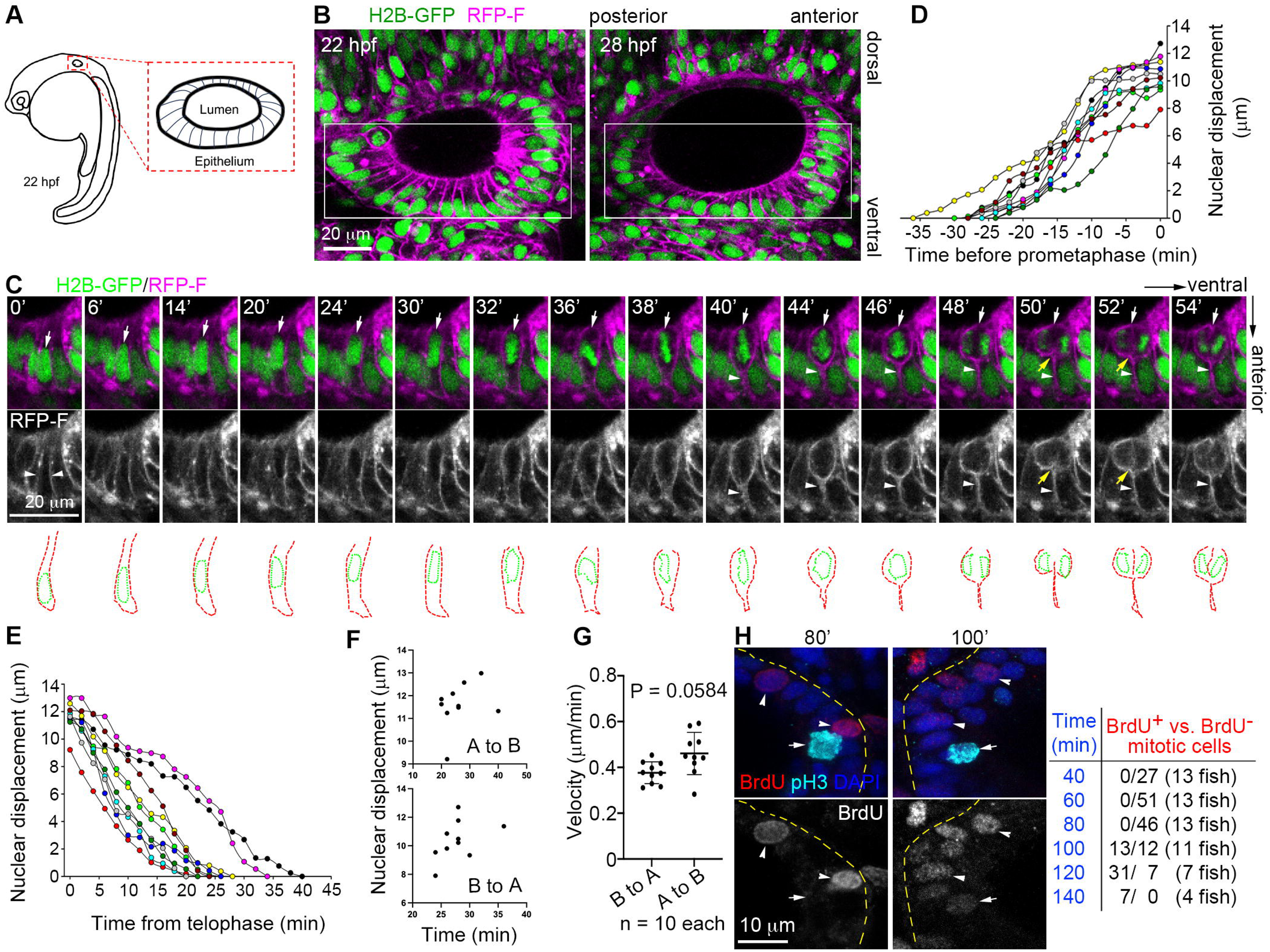
INM in zebrafish otic vesicle epithelium begins in late G2. **(A)** Schematic of the zebrafish otic vesicle at approximately 22 hpf. **(B)** Morphology of the otic vesicle epithelium at 22 hpf and 28 hpf. Embryos were microinjected with mRNA to express H2B-GFP (chromatin, green) and RFP-F (cell membrane, magenta). The boxed regions indicate areas used for INM analyses. Scale bar, 20 µm. **(C)** Selected frames from Supplementary Video S2. H2B-GFP (green) and RFP-F (magenta) label chromatin and the plasma membrane, respectively. Single-channel images of RFP-F are shown in the middle. White arrows indicate the nucleus or chromosomes of the tracked cell, white arrowheads highlight its basal protrusion and plasma membrane, and yellow arrows denote the cleavage furrow. The cell boundary and nucleolus of the tracked cell are outlined below. Scale bar, 20 µm. **(D, E)** Nuclear trajectories before prometaphase (D) and after telophase (E). For daughter nuclei, only one that could be clearly tracked was analyzed. n = 10 cells from 4 embryos each group. **(F)** Correlation of nuclear displacement and travel time for basal-to-apical (B to A) and apical-to-basal (A to B) migrations. n = 10 cells from 4 embryos each group. **(G)** Velocity of nuclear movements. n = 10 cells from 4 embryos each group. Nested t test. **(H)** Estimation of G2 length. Embryos at 24 hpf were labeled with BrdU, fixed at the indicated time points, and stained to visualize BrdU (red), pH3 (cyan), and DAPI (blue). Representative S-phase cells (arrowheads) and mitotic cells (pH3-positive, arrows) are shown. Yellow dashed lines indicate the basal membrane. Scale bar, 10 µm.

We next quantified nuclear dynamics relative to mitotic progression. Prophase-prometaphase and anaphase-telophase transitions were readily identifiable in time-lapse sequences (Fig. 1C; Supplementary Videos S1, S2), enabling precise timing of apical and basal nuclear movement (Fig. 1D-G). Apical migration began 27.8 ± 3.5 min before prometaphase onset, with an average velocity of 0.38 ± 0.05 µm/min (Fig. 1D, F, and G). Following cytokinesis, nuclei returned basally over 26.2 ± 6.4 min, moving at 0.46 ± 0.09 µm/min on average (Fig. 1E, F, and G). Migration time shows a trend toward correlation with nuclear migration distance (or displacement; Fig. 1F).

Apically directed INM is typically associated with G2 entry^10, 11, 16, 19^. To determine where INM initiation lies within G2 in this system, we measured G2 duration using BrdU pulse-labeling. Embryos were exposed to BrdU at 24 hpf and fixed at sequential time points. Mitotic cells were identified by pH3 immunostaining^36^. No BrdU^+^ mitoses were detected at 40-, 60-, or 80-minutes post-labeling (Fig. 1H). In contrast, BrdU^+^ mitotic cells appeared consistently at 100-140 minutes, indicating a G2 period of ∼80-100 min. Given that apical migration begins ∼28 min before mitosis (Fig. 1D), we conclude that INM initiates in late G2 in the zebrafish otic vesicle epithelium.

### INM in the mouse epididymis resembles that in zebrafish otic vesicle

To test whether the dynamics observed in zebrafish represent conserved epithelial behavior, we examined the murine epididymis, a tubular organ lined by simple columnar epithelial cells. While the spatial distribution of mitotic activity along the epididymal tubule has been previously reported, polarized epithelial division in this tissue has not been characterized (Fig. 2A)^7, 37^. Examination of cryosections stained for chromatin and pH3 revealed that mitotic events peaked around five weeks postpartum and localized predominantly to the initial segment (Fig. 2A-B). G2-phase nuclei, identified by punctate pH3-positive foci^36^, were detected in both basal and middle regions (Fig. 2C, panels 1, 3, 4), whereas prophase cells with condensed chromosomes and strong pH3 signal were restricted to the middle-apical domain (Fig. 2C, panel 2). Prometaphase, metaphase, and anaphase chromosomes clustered apically, whereas telophase nuclei reappeared in the middle region, consistent with a return migration (Fig. 2C, panels 2-4). Mitotic cells divided in the planar orientation at the apical surface, supporting that epididymal epithelial cells undergo INM followed by apical mitosis, similar to zebrafish otic epithelium.

**Figure 2.**
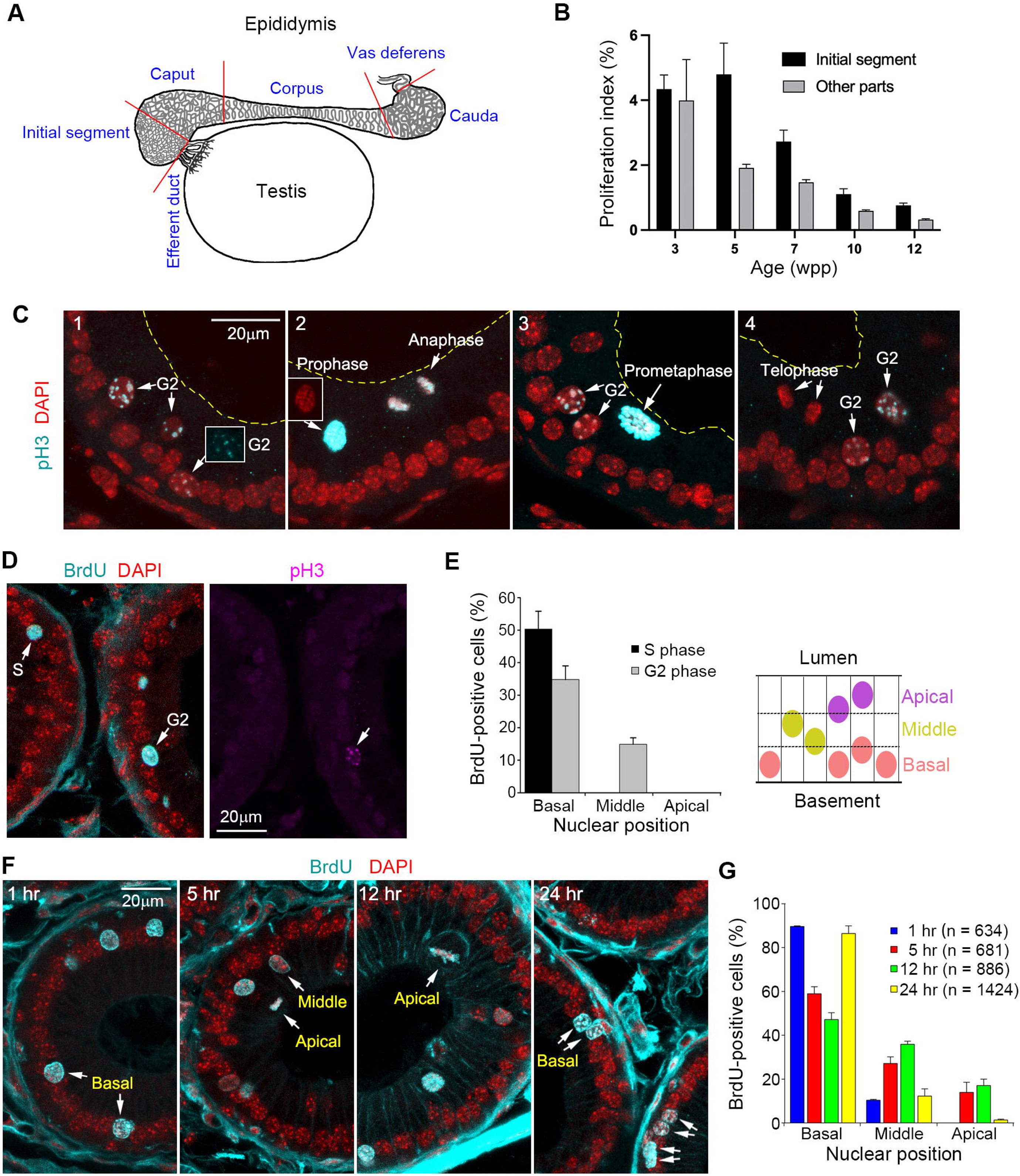
INM in the mouse epididymis resembles that in the zebrafish otic vesicle. **(A)** Schematic of the anatomical structure of the mouse epididymis. **(B)** Quantification of mitotic events. Mitotic cells were identified based on chromosome morphology and/or strong pH3 staining (see C). Three replicates per group were analyzed. **(C)** Section images showing nuclear positioning at different cell cycle stages. Insets display pH3 (cyan) or DAPI (red) staining of the indicated nuclei. Yellow lines mark the apical surface. Scale bar, 20 µm. **(D, E)** Section images and quantification (E) showing S-phase and G2-phase nuclei 1 hour after BrdU (cyan) labeling. Nuclear position is defined by the location of the majority of the nuclear area relative to the height of the epithelium. n = 750 from 5 epididymides. Scale bar, 20 µm. **(F, G)** Section images and quantification showing the distribution of BrdU (cyan)-positive nuclei or chromosome clusters at the indicated time points. Arrows in (F) indicate representative nuclei or chromosome clusters. Quantification from 4 epididymides for each time point. Scale bar, 20 µm.

To refine the timing of basal-to-apical migration, we administered BrdU intraperitoneally and analyzed tissue one hour later. Among 750 BrdU^+^ cells examined, all S-phase nuclei (BrdU^+^ pH3^-^) resided basally (Fig. 2D-E). Most G2-phase nuclei (BrdU^+^ pH3^+^) were also basal, with a minority shifted to the middle region, indicating that apical translocation begins during late G2 rather than immediately upon G2 entry.

We next traced nuclear position over time using a BrdU pulse-chase. At one hour, 89.6% of BrdU^+^ nuclei were basal. By 5-12 hours, labeled nuclei were distributed throughout the epithelium (Fig. 2F-G). At 24 hours, BrdU^+^ nuclei again peaked basally (86.4%), with frequent paired nuclei indicating daughter cell residency post-division (Fig. 2F-G). Together, these data demonstrate that mouse epididymal epithelial cells exhibit basal S-phase positioning, late-G2 apical nuclear migration, apical planar division, and post-mitotic basal return, with timing and cellular features similar to those observed in the zebrafish otic vesicle epithelium.

### Cell shape dynamics and basal lamina attachment during INM and mitosis

To gain insight into the mechanisms underlying INM, we examined how cell shape changes during nuclear migration and mitosis. In the mouse epididymis, visualization of lateral adherens junctions using E-cadherin staining or cortical F-actin (visualized by phalloidin staining^38, 39^) revealed that columnar morphology was maintained from interphase through prophase (Fig. 3A-C). As cells entered prometaphase, they began rounding while retaining a thin basal projection attached to the basal lamina, separating the main cell body from the basal membrane (Fig. 3A and C, arrowheads), which persisted through division (Fig. 3A). Intensity of cortical F-actin signals (versus non-dividing neighbors) increased in prometaphase, peaked in metaphase, and decreased after division (Fig. 3A, B). Although E-cadherin intensity at lateral membranes remained relatively constant (Fig. 3C, and F), phospho-E-cadherin and β-catenin signals increased markedly in prometaphase and metaphase cells (Fig. 3D-F). β-catenin staining intensity in metaphase cells is ∼2.044-fold higher than in neighboring non-dividing cells (Fig. 3E). These findings suggest more stable lateral adhesion between mitotic cells and their interphase neighbors^38–40^.

**Figure 3.**
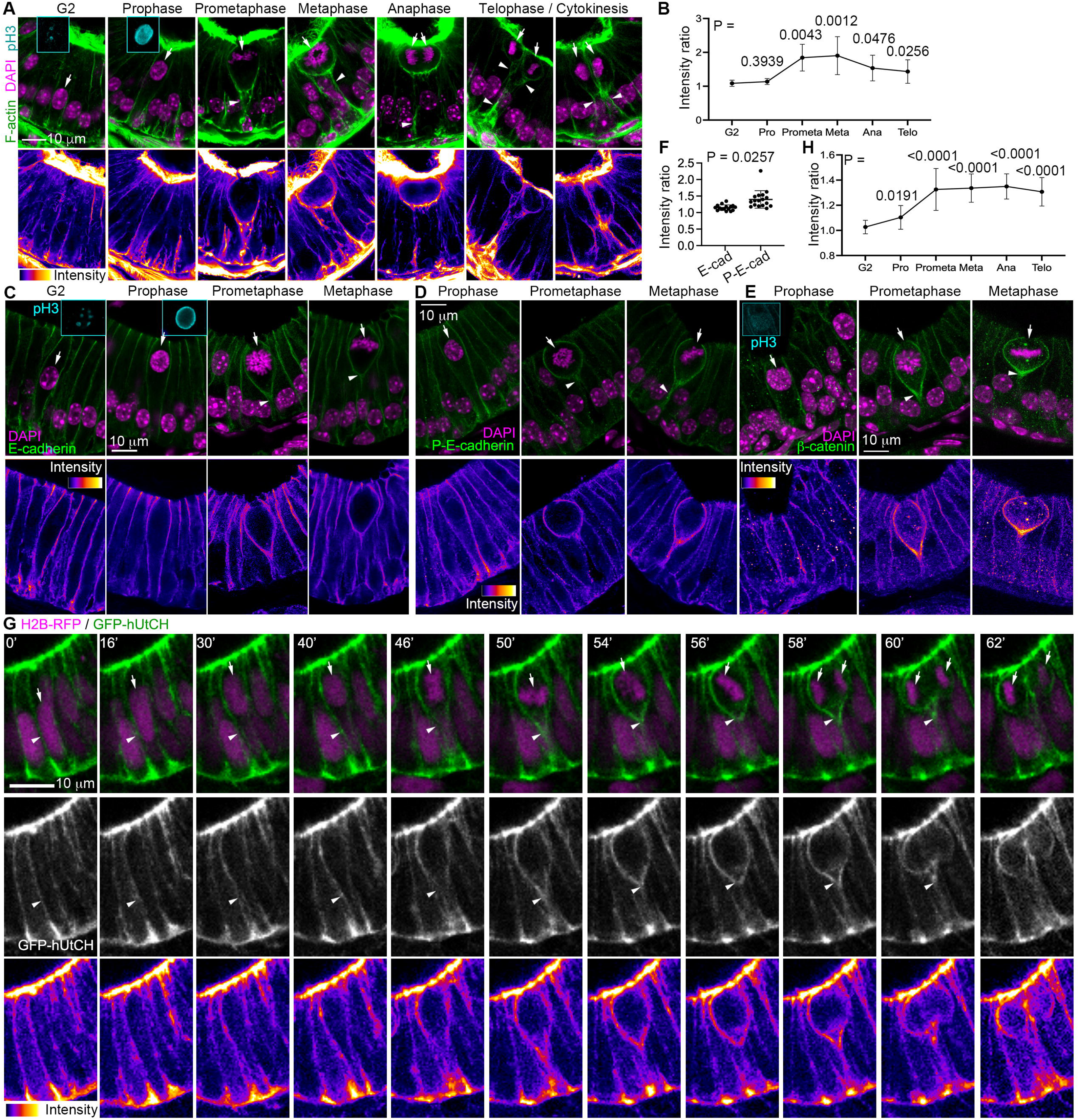
Cell shape dynamics and basal lamina attachment during INM and mitosis. **(A)** Section images of mouse epididymis epithelium showing F-actin staining (green). Insets show pH3 staining (cyan) in G2 or prophase cells. Arrows indicate G2 or M-phase cells, and arrowheads point to the narrow basal protrusions of mitotic cells. DAPI staining is shown in magenta. Fluorescent intensity of F-action signals in is shown at the bottom. Scale bars, 10 µm. **(B)** Quantification of the F-actin intensity ratio (INM or dividing cells versus non-migrating and non-dividing neighboring cells) across cell-cycle stages. F-actin intensity was measured at the lateral membranes of non-migrating/non-dividing cells and at both the lateral and basal side membrane of rounded dividing cells (excluding the basal stalk). Two-tailed Mann-Whitney U test was performed relative to the G2 phase. n = 36 cells from 3 experiments. **(C-E)** Section images of mouse epididymis epithelium showing cell-cell interfaces labeled with E-cadherin (C), phospho-E-cadherin (P-E-cadherin, D) and β-catenin (E) in green. Insets show pH3 staining (cyan) in G2 or prophase cells. Arrows indicate G2 or M-phase cells, and arrowheads point to the narrow basal protrusions of mitotic cells. DAPI staining is shown in magenta. Fluorescent intensity of E-cadherin, P-E-cadherin, or β-catenin signals is shown at the bottom. Scale bars, 10 µm. **(F)** Quantification of E-cadherin (E-cad) or phosphor-E-cadherin (P-E-cad) intensity ratio (metaphase cells versus non-dividing neighboring cells). Signal intensity was measured at the lateral membranes of non-dividing cells and at both the lateral and basal side membrane of rounded dividing cells (excluding the basal stalk). n = 16 cells from 4 samples for E-cad and 17 cells from 4 samples for P-E-cad. Nested t-test. **(G)** Selected frames of zebrafish otic vesicle epithelial cells from Video S3 showing GFP-hUtCH (green) and H2B-RFP (magenta). Single-channel GFP-hUtCH images are shown in grayscale at the bottom. Arrows indicate the nucleus or chromosomes of the tracked cell, while arrowheads highlight its basal process and plasma membrane. Fluorescent intensity of GFP signals is shown at the bottom. Scale bar, 10 µm. **(H)** Quantification of the GFP-hUtCH intensity ratio (INM or dividing cells versus non-migrating and non-dividing neighboring cells) across cell-cycle stages. F-actin intensity was measured at the lateral membranes of non-migrating/non-dividing cells and at both the lateral and basal side membrane of rounded dividing cells (excluding the basal stalk). Two-tailed Mann-Whitney U test was performed relative to the G2 phase. n = 19 traced cells from 3 embryos. Each cell has multiple stages quantified.

We next asked whether similar dynamics occur in zebrafish. Live imaging of otic vesicle cells expressing H2B-GFP and RFP-F showed that columnar shape was largely preserved during basal-to-apical INM, consistent with findings in the epididymis (Fig. 1C, Supplementary Video S2). Beginning at prometaphase, the basal portion of the lateral cortex gradually constricted, leaving a stalk-like basal attachment as the cell rounded (Fig. 1C, 32-48’, arrowheads). Cleavage furrow ingression initiated basally (Fig. 1C, 50’, yellow arrow) and reached the apical surface within ∼2 minutes (Fig. 1C, 52’). To further characterize actin behavior, we expressed GFP-hUtrCH, a probe preferentially labeling stable cortical F-actin^26, 41^. GFP-hUtrCH localized to the cell cortex in otic epithelium (Fig. 3G, Supplementary Video S3), confirming effective labeling. Live imaging revealed timing and morphological features similar to those described above, including prometaphase rounding (30’-56’; Fig. 3G), persistent basal lamina attachment (50’-62’), and basal-to-apical cleavage furrow progression (58’-62’), which initiates from the basal process (Fig. 3G). As expected, increased GFP-hUtrCH intensity during rounding and furrow ingression (Fig. 3G, bottom) further demonstrate actomyosin involvement and potential cortex stabilization. We observed comparable hUtrCH intensity dynamics across cell-cycle stages in zebrafish (Fig. 3H) relative to the mouse epithelium (Fig. 3B). Together, these results show that columnar epithelial cells in both mouse epididymis and zebrafish otic vesicle undergo prometaphase cell rounding while preserving a basal anchoring process. Our data suggest that cytokinesis initiates from the basal side in the zebrafish otic epithelium; however, the rapidly evolving dynamics of cytokinesis could not be captured in fixed mouse tissues. These behaviors align with prior findings in *Drosophila* optic lobe, zebrafish hindbrain, and rodent neocortex^12, 42^, suggesting a conserved mitotic program among columnar epithelia and indicating that the zebrafish otic vesicle epithelium and mouse epididymal epithelium are well-suited models for addressing related questions.

### Myosin II regulates mitotic cell rounding, spindle orientation, and epithelial Integrity, but is dispensable for basal-to-apical migration

Our live imaging of F-actin dynamics suggested that actomyosin contractility contributes to division behaviors in columnar epithelia. To test the requirement for myosin II-mediated contractility, we perturbed myosin II function in zebrafish otic vesicle epithelium using blebbistatin, a well-established inhibitor of non-muscle myosin II^43^. Myosin II is known to generate cortical tension in non-muscle cells and has been linked to INM in pseudostratified neuroepithelia^25, 26^. As expected, blebbistatin treatment did not disrupt actin polymerization, as GFP-hUtrCH localization remained comparable to controls (Fig. 4A; Supplementary Videos S4, S5). Inhibition was effective as embryos displayed pronounced cytokinesis defects (Fig. 4A; Supplementary Videos S4, S5)^43^. Strikingly, blebbistatin treatment had no effect on basal-to-apical INM. Both the total number of migration events per otic vesicle and the average velocity (0.36 ± 0.08 µm/min) for basal-to-apical migration were indistinguishable from vehicle controls (0.39 ± 0.07 µm/min) (Fig. 4B-C) within the imaging window. Similarly, no significant difference in nuclear migration distance or time was observed (Fig. 4D, E). These results suggest that apical INM is myosin II-independent in this simple columnar epithelium. This contrasts with the pseudostratified fly neuroepithelium, where INM is driven by myosin-dependent mitotic rounding^12^.

**Figure 4.**
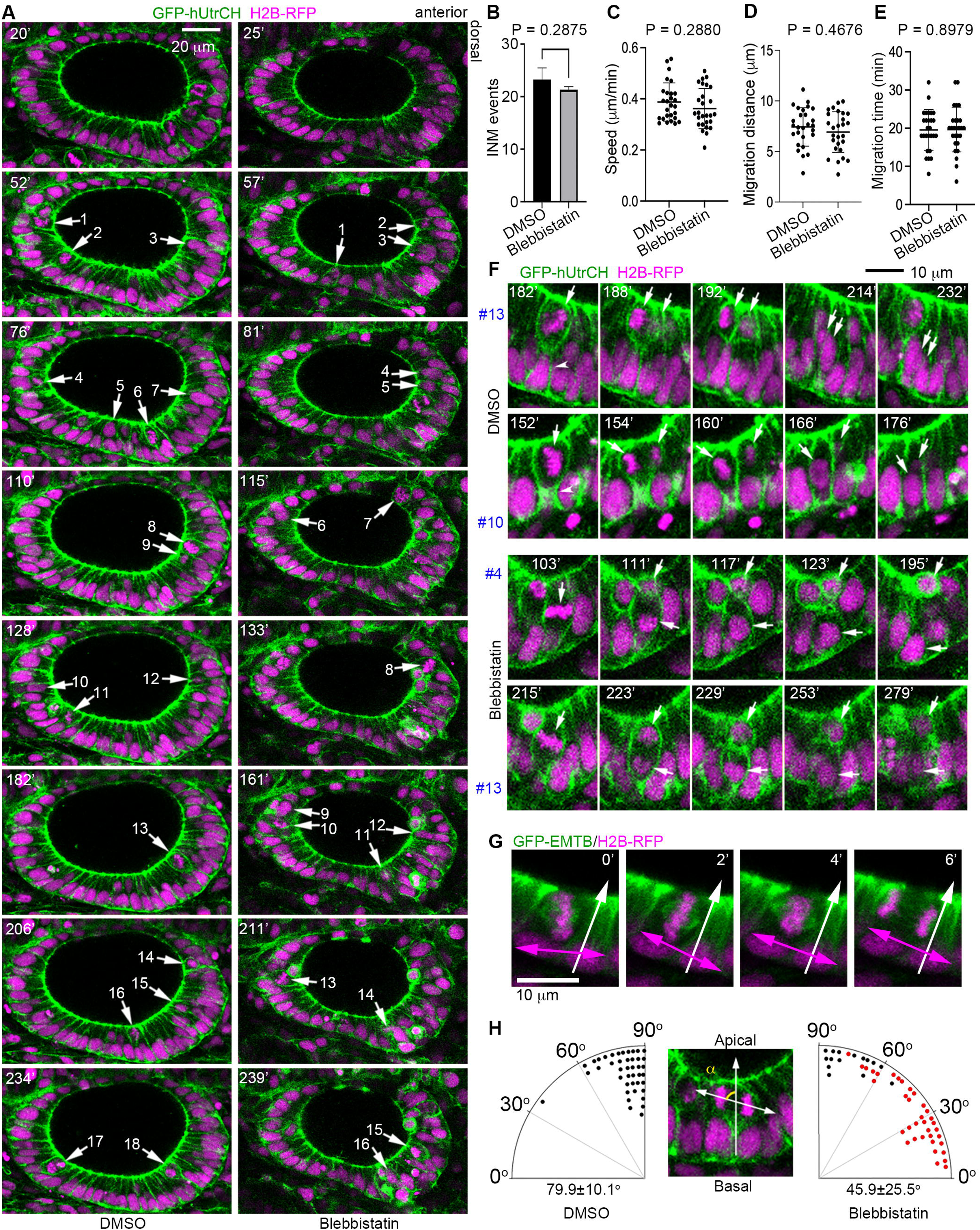
Myosin II activity regulates mitotic cell rounding, spindle orientation, and epithelial Integrity in zebrafish otic vesicle epithelia. **(A)** Selected frames from Supplementary Videos S4 and S5 showing INM events. H2B-RFP (magenta) and GFP-hUtrCH (green) are shown. Time is indicated relative to the addition of DMSO or blebbistatin. Each number represents a nuclear migration event and marks a mitotic cell. Scale bar, 20 µm. **(B)** Quantification of INM events during a 230-min window while epithelial cells remained columnar, starting 20 or 25 min after drug addition. Two-tailed Mann-Whitney U test. n = 4 (DMSO) or 3 (Blebbistatin) fish. **(C-E)** Quantification of migration speed (C), distance (D), and time (E) of apically directed INM. n = 27 cells from 3 fish each group. Nested t tests. **(F)** Representative images from Supplementary Videos S4 and S5 showing mitotic rounding, planar division, and daughter cell insertion after treatment. The locations of mitotic cells in the otic vesicle are indicated in (A) by the corresponding numbers. Arrows indicate parental or daughter nuclei. Arrowheads denote the basal process. Scale bar, 10 µm. **(G)** Representative frames showing spindle orientation with GFP-EMTB (green) and H2B-RFP (magenta). Magenta arrows indicate spindle orientations, whereas white arrows denote apical-basal polarity. Scale bar, 10 µm. **(H)** Statistical analysis of cell division angles. Each mitotic cell is represented as a dot in the radial histogram. Cells with apically retained daughter nuclei are highlighted in red. n = 42 (DMSO) or 51 (Blebbistatin) cells from 3 fish per group.

Despite normal INM, several mitotic defects emerged upon myosin inhibition. First, blebbistatin-treated cells failed to round properly: 98.3% of control cells (58/59) underwent robust rounding at prometaphase, while 73.1% of treated cells (38/52) retained a diamond-shaped morphology (maintaining a wide basal connection) through prometaphase and metaphase (Fig. 4F; Supplementary Videos S4, S5). Second, although division remained apical, spindle orientation was frequently misaligned. Whereas 97.6% of control cells (41/42) divided planarly (60°-90° relative to the apical surface), only 39.2% of treated cells (20/51) maintained this orientation; an equal fraction (39.2%) divided nearly along the apical-basal axis (0°-30°) (Fig. 4F-H). Third, daughter cells often failed to reintegrate into the epithelium. In controls, both post-mitotic nuclei returned basally within ∼30 min of anaphase onset (Fig. 4A, F; Supplementary Video S4). By contrast, 70.6% of blebbistatin-treated divisions (36/51) retained one nucleus apically for >50 min (Fig. 4A, F; Supplementary Video S5). Of these, 33 remained apical for the duration of imaging (Fig. 4H, red dots). Retained cells remained rounded with sustained GFP-hUtrCH enrichment, suggesting defective reinsertion into the epithelium. Notably, nuclear retention strongly correlated with division angle: 96.8% of divisions at 0°-60° (30/31) resulted in apical retention, compared to only 30% (6/20) at 60°-90° (Fig. 4H). Cells dividing closer to the lumen were most prone to misintegration (Fig. 4F). Because blebbistatin affects all cells, whether non-cell-autonomous effects on mechanical force within the tissue contribute to the observed spindle orientation defect warrants further investigation.

### Basal-to-apical INM requires microtubules

Both MTs and F-actin have been implicated in regulating INM^10, 11, 19^. To define the specific contribution of MTs in our INM models, we first performed immunostaining against α-tubulin in mouse epididymal epithelium, revealing MT bundles aligned along the apicobasal axis (Fig. 5A). Antibody staining for acetylated tubulin (Ace-tub) revealed a similar distribution for stable microtubules (Fig. 5B, C). Consistently, live imaging of the zebrafish otic vesicle epithelium using EMTB-GFP, a fluorescent MT reporter^44–46^, suggested a similar pattern of MTs aligned along the apicobasal axis (Fig. 5D; Supplementary Video S6).

**Figure 5.**
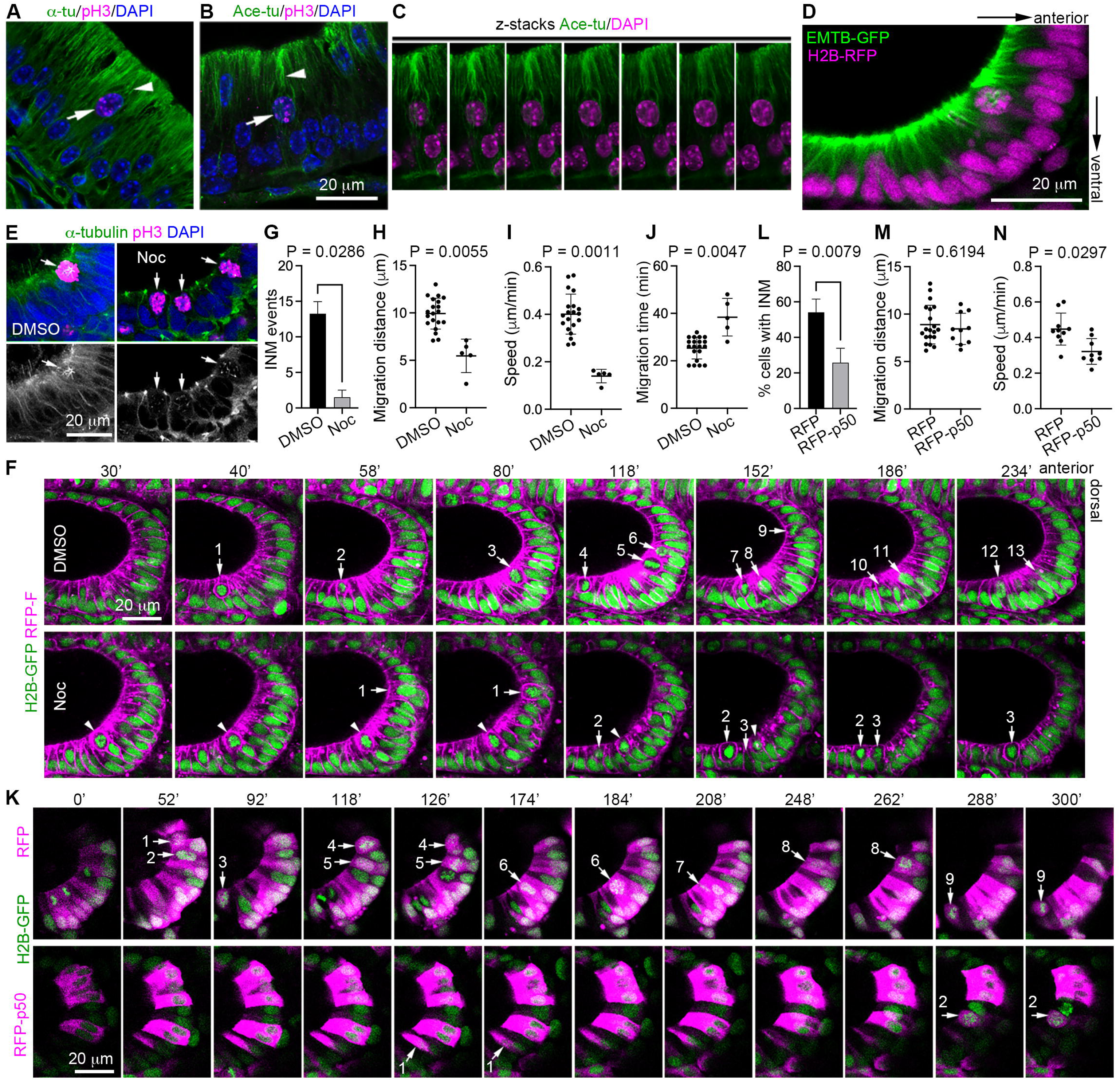
INM in zebrafish otic vesicle requires MTs and dynein. **(A, B)** Section images of epididymal epithelium stained for α-tubulin (A, green), acetylated tubulin (Ace-tu, B, green), pH3 (magenta), and DAPI (blue). Arrows indicate pH3^+^ cells, and arrowheads highlight microtubule bundles. Scale bars, 20 µm. **(C)** Z-stack images of the same cell as in (B), showing Ace-tu (green) and DAPI (magenta) staining. **(D)** Video frame showing EMTB-GFP (green) and H2B-RFP (magenta). Scale bar, 20 µm. **(E)** Nocodazole-induced microtubule disassembly. Zebrafish otic vesicle was stained for α-tubulin (green) and pH3 (magenta). DAPI (blue) staining is also shown. Single-channel α-tubulin images are displayed in grayscale at the bottom. Scale bar, 20 µm. **(F)** Images from Supplementary Videos S7 and S8 showing INM events (arrows) in the otic vesicle after nocodazole or vehicle (DMSO) treatment. H2B-GFP (green) and RFP-F (magenta) are shown. Time is indicated relative to drug addition. The arrowhead marks a cell arrested in M phase from the beginning of imaging. Scale bar, 20 µm. **(G)** Quantification of INM events during a 206-min window, starting 30 min after drug addition. Data from three embryos per condition. Two-tailed Mann-Whitney U test. n = 3 fish each group. **(H)** Quantification of migration distance of apically directed INM. n = 20 cells from 3 fish for the DMSO group and 5 cells from 4 fish for the nocodazole group. Nested t test. **(I)** Quantification of migration speed. n = 20 cells from 3 fish for the DMSO group and 5 cells from 4 fish for the nocodazole group. Nested t test. **(J)** Quantification of migration time. n = 20 cells from 3 fish for the DMSO group and 5 cells from 4 fish for the nocodazole group. Nested t test. **(K)** Images from Supplementary Videos S9 and S10 showing INM events (arrows) in the otic vesicle. RFP or RFP-p50 was overexpressed under the hsp70 promoter (magenta). H2B-GFP was expressed to label chromatin (green). Each number represents an INM event in RFP-positive cells. Scale bar, 20 µm. **(L)** Quantification of INM events over 300 min of imaging. Data from 5 embryos per condition with 61 cells (RFP-C1) and 53 cells (RFP-P50) analyzed, respectively. Two-tailed Mann-Whitney U test. **(M)** Quantification of migration distance of apically directed INM. n = 20 cells from 5 fish for the RFP group and 10 cells from 5 fish for the RFP-P50 group. Nested t test. **(N)** Quantification of migration speed of apically directed INM. n = 10 cells from 5 fish for the RFP group and 9 cells from 4 fish for the RFP-P50 group. Nested t test.

To functionally assess MT requirements, we treated zebrafish embryos with 0.5 µg/ml nocodazole to disrupt MT polymerization. Drug efficacy was confirmed by fixation 120 min after treatment, which showed absent or fragmented mitotic spindles in mitotic cells of the otic vesicle, whereas vehicle-treated embryos displayed normal bipolar spindles (Fig. 5E, arrows). Interphase MT networks were also largely lost following nocodazole treatment (Fig. 5E). Live imaging initiated ∼20 min after drug addition showed that many mitotic cells arrested in M-phase (Fig. 5F, arrowhead), confirming effective MT inhibition. Time-lapse imaging further revealed a pronounced reduction in INM events per otic vesicle (Fig. 5F, G; Supplementary Videos S7, S8). Because nocodazole progressively thinned the epithelium, quantitative analysis was confined to the thicker anterior-ventral region between 30-236 min post-treatment, when epithelial morphology remained columnar (Fig. 5F; Supplementary Video S8). Relative to vehicle controls, nocodazole caused an average 7.8-fold reduction in the number of INM event within a 206-min imaging window (Fig. 5G). We observed only 6 INM events from four nocodazole-treated embryos, compared with 53 INM events from four control embryos. For this limited number of basal-to-apical migration, both migration distance and migration speed were significantly reduced, while migration time was increased upon nocodazole treatment (Fig. 5H-J). Together, these results indicate that microtubule depolymerization impairs both INM initiation and migration speed, suggesting that microtubules play an essential role in basal-to-apical INM in vertebrate columnar epithelia.

### Cytoplasmic dynein activity is critical for basal-to-apical INM

Rodent neuroepithelia studies suggest that cytoplasmic dynein, a microtubule-based, minus-end-directed motor, is a key driver of basal-to-apical nuclear movement. This activity is supported by proteins located both at the apically anchored centrosome (e.g., Cep120, TACCs, Hook3, PCM1), which maintain polarized microtubule arrays^20–23^, and at the nuclear envelope (e.g., Syne1/2, Sun1/2), which link nuclear membranes to dynein^10, 24^. To test dynein involvement in zebrafish INM, we overexpressed RFP-tagged human p50, a dominant inhibitor of dynein function^47–49^. To avoid early developmental defects due to dynein disruption, RFP-p50 was placed under the inducible *hsp70* promoter and embryos were heat-shocked at 20 hpf for 1 hr to trigger expression^50^. Live imaging was initiated ∼1.5 hr later, when robust RFP signal became visible. Compared with RFP controls, RFP-p50 overexpression markedly reduced apically directed nuclear migration events (Fig. 5K and L; Supplementary Videos S9, S10). Over a 300-min imaging window, 55 ± 10% of RFP-positive control cells underwent apical INM, whereas only 21 ± 5% of RFP-p50-expressing cells did so (Fig. 5L). The migration distance and speed were comparable between the two groups (Fig. 5M and N). These results demonstrate that cytoplasmic dynein is essential for MT-dependent basal-to-apical INM in zebrafish epithelia.

### Nuclear migration is required for M phase entry

Notably, in long columnar cells treated with nocodazole, mitotic entry (defined by the onset of H2B-GFP signal condensation into distinct, bright structures) occurred only after successful apical migration of the nucleus (Fig. 5F; Supplementary Video S8). In contrast, in the naturally thinner posterior region of the otic epithelium, where nuclei remain near the apical surface, cells frequently entered mitosis even in the presence of nocodazole (Supplementary Video S8), suggesting that nuclear position, rather than MT polymerization *per se*, is coupled to M-phase onset. To probe this relationship further, immunostaining of mouse epididymal epithelium showed that centrosomes occupy apical positions throughout INM, and that centriole separation occurred only once nuclei reached the apical surface (Fig. 6A). Live EMTB-GFP imaging confirmed similar MTOC behavior in the zebrafish otic vesicle epithelium (Fig. 6B; Supplementary Video S6). We next examined cyclin B1, a late-G2/M-phase marker. In migrating nuclei, cyclin B1 remained cytoplasmic and translocated into the nucleus only after apical arrival (Fig. 6C, D). Notably, after long-term nocodazole treatment (up to 8 h) in the zebrafish otic epithelium, most cells entered M phase when the epithelium became thin and nuclei were positioned near the apical membrane (Supplementary Fig. S1). Consistent with previous studies^51, 52^, these data suggest that apical nuclear positioning is tightly linked to mitotic commitment, and that reaching centrosomes at the apical surface is likely a prerequisite for M-phase entry.

**Figure 6.**
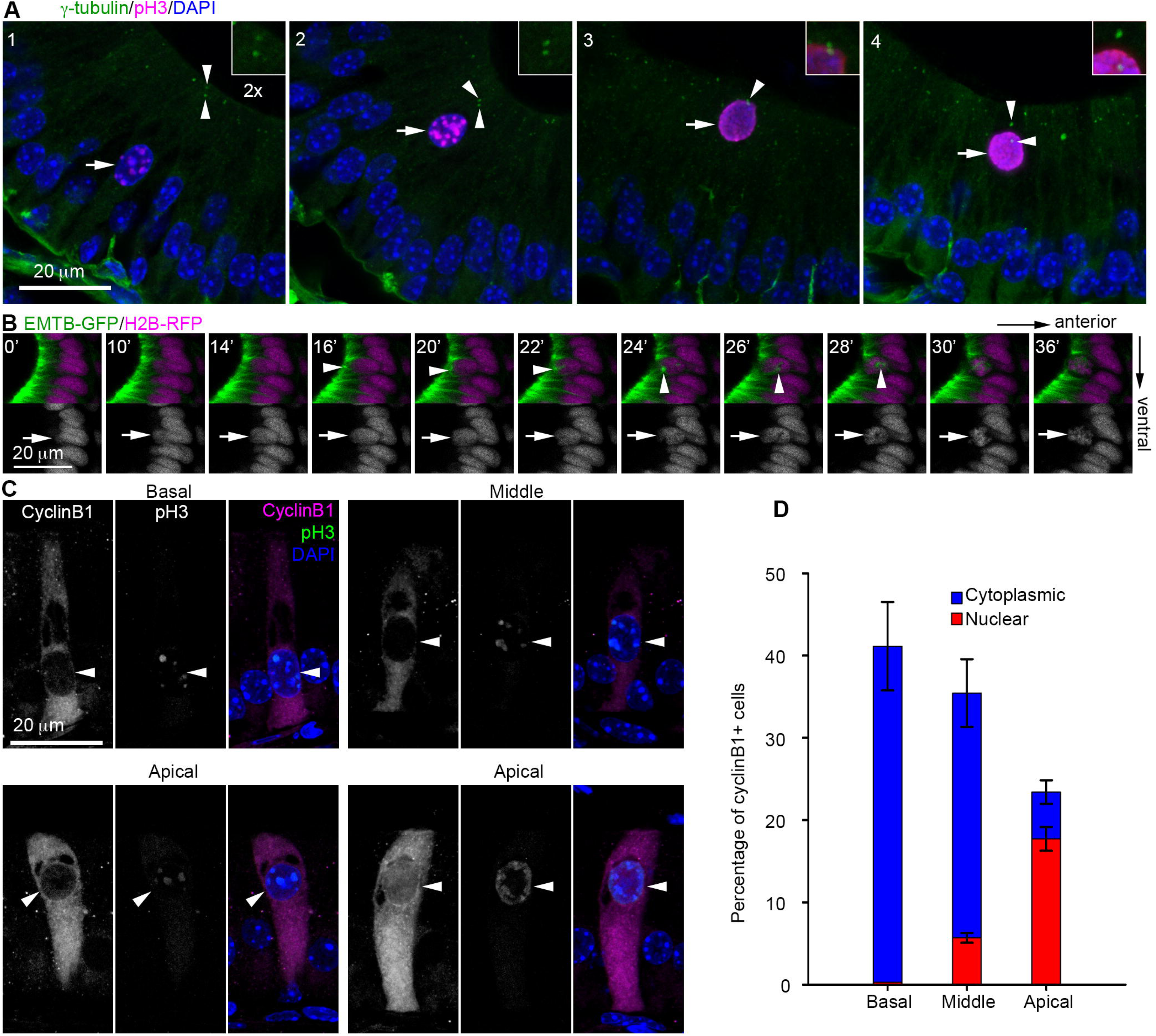
Nuclear migration is essential for M phase onset. **(A)** Sections of mouse epididymal epithelium stained for γ-tubulin (green), pH3 (magenta), and DAPI (blue). Centrosomes (arrowheads) localize near the apical surface during nuclear migration, while centrioles move to opposite sides of the nucleus (arrows) only after it reaches the apical side. **(B)** Frames from Supplementary Video S6 of zebrafish otic vesicle epithelial cells. Nuclei labeled with H2B-RFP (magenta) and stable microtubules labeled with EMTB-GFP (green) show MTOC/centrosomes (arrowheads) migrating to opposite sides of the nucleus (arrows) after apical migration to form the spindle. **(C)** Sections of mouse epididymal epithelium stained for cyclin B1 (magenta), pH3 (green), and DAPI (blue). Scale bar, 20 µm. **(D)** Quantification for images shown in (C). Subcellular localization of Cyclin B1 grouped by position of nuclei in the basal, middle and apical parts of the cell. Blue = cytoplasmic, red = nuclear Cyclin B1 expression. n = 299 cells from three mice.

## DISCUSSION

Previous work has described the morphogenesis of the zebrafish otic vesicle^31–33^. However, here we adapted this system as a powerful live, physiological, and genetically tractable model for real-time visualization of epithelial nuclear dynamics and mitosis. This approach overcomes key limitations of fixed-tissue methods, enabling direct observation of nuclear behaviors throughout the cell cycle. By extending our analysis to the mouse epididymal epithelium, we further uncovered shared cellular features of INM in vertebrate columnar epithelia.

Our findings indicate that basal-to-apical INM initiates during late G2 in the zebrafish otic vesicle epithelium, and similarly in the mouse epididymis, where nuclear translocation predominantly occurs after G2 entry (Fig. 7). Interestingly, while nuclear oscillations are frequently observed in neuroepithelia, a rapid, directional basal-to-apical movement occurs just prior to mitosis^20, 26, 53^. This movement appears to initiate in mid-to-late G2 across different epithelial systems. For example, directional INM lasts 5-30 minutes in the developing zebrafish retina ^26^ and approximately 60 minutes in the mouse neural tube^14, 45^. Thus, the late onset of INM during G2 may be a conserved feature across both simple columnar and pseudostratified epithelia.

**Fig. 7.**
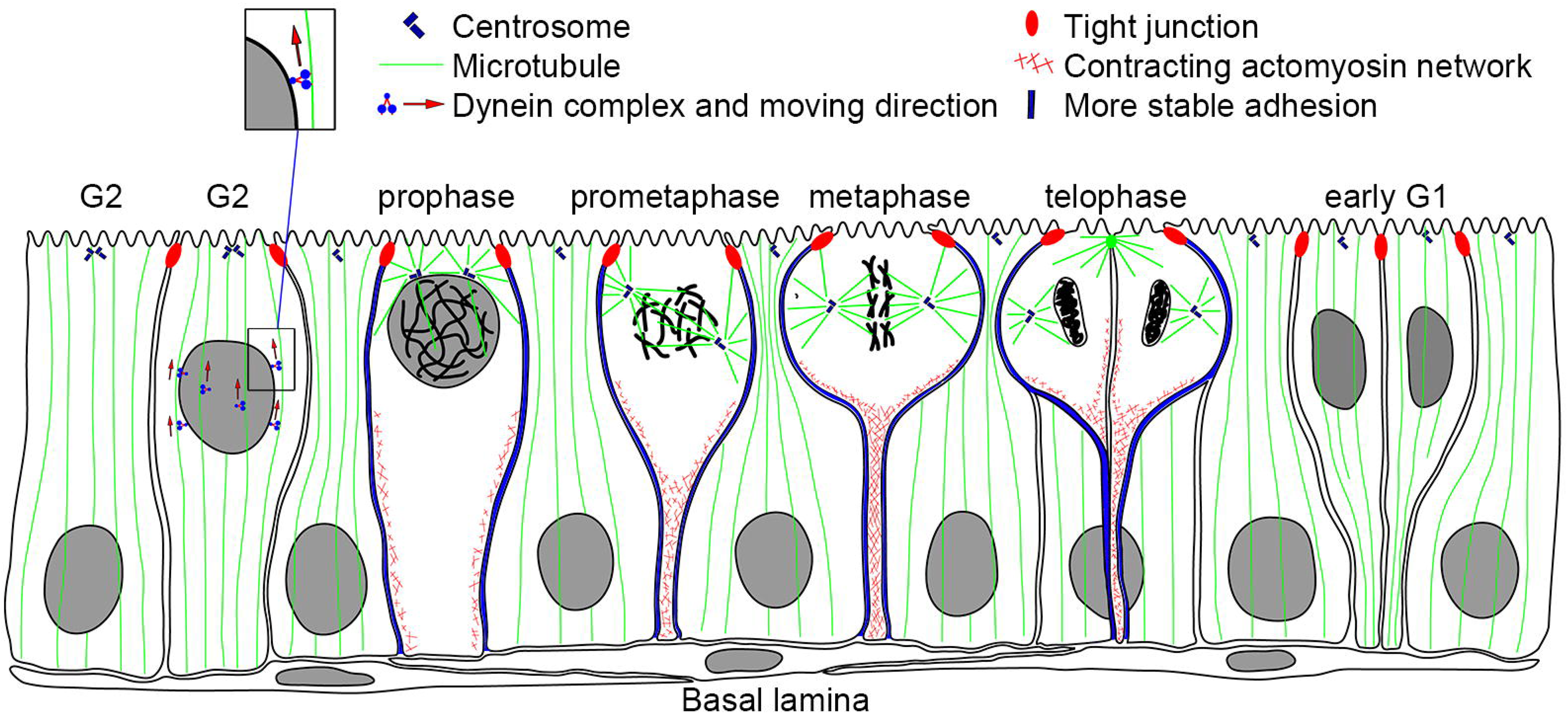
Summary of INM and cell division in simple columnar epithelium. A schematic summarizing apically directed INM, which is driven by dynein in late G2. During prometaphase, actomyosin mediates mitotic rounding by constricting the basolateral membrane, while lateral adhesions to neighboring interphase cells are strengthened. Mitotic cells maintain attachment to the basal lamina during planar division.

Mechanistically, we show that basal-to-apical INM in the otic vesicle requires MTs and dynein (Fig. 7), whereas inhibition of myosin II activity had minimal effects on this process. This diverges from neuroepithelia, where MTs, dynein, and actomyosin contractility all contribute to nuclear movement^10, 13, 19^. A likely explanation lies in differences in epithelial architecture. In pseudostratified tissues, nuclei migrate through narrow, elongated processes, necessitating both MT-based pulling and actomyosin-driven squeezing forces along lateral membranes^10, 13, 19^. In contrast, columnar epithelia maintain the columnar shape before the mitotic rounding, and actomyosin forces largely dispensable for basal-to-apical migration. Because dynein inhibition or MT disassembly abolished apical migration, we could not determine whether MT-driven forces also contribute to basally directed nuclear movement. Although blebbistatin-treated embryos showed defective planar division, one or both daughter nuclei still returned to the basal side, suggesting that myosin II is not strictly required for apical-to-basal nuclear migration.

We further show that mitotic cell rounding in both zebrafish otic vesicle and mouse epididymal epithelium occurs during prometaphase (Fig. 7). The maintenance of basal adhesion during mitosis, as observed in our study, has also been reported in other epithelia, including the *Drosophila* optic lobe, zebrafish hindbrain, rodent neocortex, and mouse intestine and kidney, suggesting that this feature is conserved across both simple columnar and pseudostratified epithelia^6, 10–12, 19, 42, 54^. Our results identify myosin II as a primary driver of mitotic rounding. The increased levels of F-actin, phospho-E-cadherin, and β-catenin at lateral membranes likely reinforce actomyosin contractility, enabling lateral membrane constriction while preserving cell-cell adhesion to neighboring interphase cells (Fig. 7)^38, 39, 55^. How mitotic rounding and INM are coordinated with specific cell cycle stages remains an important open question. We also find that planar division requires myosin II activity, likely because mitotic rounding is necessary for proper spindle orientation. Given that spindle alignment tends to follow the long axis of cells and depends on cortical anchoring cues^29, 56–58^, mitotic rounding may generate the geometric and mechanical conditions needed for planar spindle orientation and successful epithelial division^3, 29, 59^.

A functional consequence of disrupted mitotic rounding becomes apparent in blebbistatin-treated zebrafish embryos. When divisions occur perpendicular to the epithelial plane (0-30°), one daughter nucleus frequently remains apical near the lumen, whereas its sister typically reinserts into the basal region. By contrast, planar divisions (60-90°) are far less likely to produce apical retention. One possible explanation is that planar division preserves both apical tight junctions^54^ and basal lamina attachment, maintaining basal-apical polarity of daughter cells (Fig. 7). Our observations in both mice (Fig. 3A, telophase/cytokinesis) and zebrafish (Fig. 3G, 58-60 min; Fig. 4F, DMSO group) are consistent with the possibility that basal contacts are maintained in both daughter cells (i.e., the division plane may pass through the basal stalk), although this interpretation requires further experimental validation. Perpendicular division, in contrast, likely compromises basal contacts, which could hinder reinsertion and destabilize epithelial architecture. These observations underscore how division orientation feeds back into epithelial homeostasis and suggest that similar mechanisms may operate broadly across simple columnar tissues.

Comparison with neuroepithelia further highlights context-dependent requirements for INM. In the neural tube and retina, disruption of INM (via cytochalasin B, Lis1/Dynein inhibition, or genetic perturbation) does not prevent cells from entering mitosis at basal or intermediate positions^23, 60–62^. INM is therefore not strictly required for cell cycle progression in those systems, although cell-cycle arrest halts INM^63^. Apical migration in the zebrafish retinal neuroepithelia is independent of centrosome position or mitotic entry site^64^. In contrast, our findings demonstrate that mitosis in the tall otic vesicle epithelium occurs exclusively at the apical surface, and disrupting MTs or dynein dramatically reduces mitotic events without generating ectopic basal divisions. The requirement for apical positioning correlates with our observation that cyclin B nuclear entry and spindle assembly occur only when the nucleus reaches the apical domain, suggesting that proximity to apical centrosomes or associated factors may be necessary for M-phase onset. Together, these results reveal that INM regulation varies across epithelial architectures, and that nuclear positioning may function as a gatekeeper for mitotic entry in columnar epithelia. Future studies should define the apical cues that governs this spatial control of mitosis and determine how centrosome-associated components, tension landscapes, or polarity determinants contribute to M-phase licensing. In addition, functional studies in mice using genetic manipulations of dynein activity or microtubules will be important for assessing the extent to which INM mechanisms are shared across species. Elucidating these mechanisms will help clarify how INM integrates with tissue growth, repair, and epithelial morphogenesis across vertebrate systems.

### Limitations of the study

Several technical limitations should be noted. First, we relied on transient mRNA expression to label histones, F-actin, and MTs, demonstrating the feasibility of this approach but introducing potential variability and overexpression-related artifacts. Stable transgenic reporters offer a more consistent alternative. For example, the *h2afv:h2afv-mCherry*^65^ or *hsp70:H2B-RFP*^66^ line can serve as a histone marker, while *actb2:mCherry-UTRN*^67^ or *actb1:lifeact-GFP*^68^ can be used for F-actin labeling. Additionally, Fluorescent Ubiquitin-Mediated Cell Cycle Indicator (FUCCI) reporters^69–71^ provide a powerful tool for tracking cell cycle progression. Moreover, the availability of inducible transgenic and gene-manipulation systems in zebrafish will facilitate more mechanistic interrogation. Second, our live imaging was performed using an inverted confocal microscope, necessitating precise embryo orientation so that the otic vesicle lies close to the coverslip for optimal signal detection. Images were obtained through a glass-bottom dish using a 20x objective, with whole-mounted embryos in low-melting-point agarose. Upright imaging with high-NA water-immersion objectives could simplify sample mounting and improve optical access, while spinning-disk microscopy would enable higher spatiotemporal resolution. Third, the morphology of the zebrafish embryonic otic vesicle evolves rapidly, with a limited window of approximately 6 hours (22-28 hpf) during which the ventral side maintains a columnar epithelial structure. This short imaging window may restrict studies of slow genetic perturbations or long-term lineage outcomes. Lastly, global labeling of F-actin does not resolve which individual cells contribute to increased cortical actin. A heat-shock-inducible mosaic labeling approach could provide single-cell resolution to more precisely capture actin dynamics. A similar strategy could also be applied to microtubule labeling to improve visualization of microtubule behavior at the single-cell level, and to myosin manipulation (e.g., via upstream regulators such as the Rho/ROCK pathway^72^) in isolated cells to test cell autonomous effects. Despite these constraints, our systems provide a uniquely accessible view of epithelial cell division *in vivo* and offer a foundation for deeper mechanistic investigation.

## Materials and Methods

### Animals

The study is reported in accordance with ARRIVE guidelines. All animal procedures were conducted in accordance with protocols approved by the Institutional Animal Care and Use Committee (IACUC). All methods were performed in accordance with relevant guidelines and regulations. Zebrafish (Ekkwill (EK) and AB strains) were raised under standard conditions and staged in hours post-fertilization (hpf) as previously described^73,74^. Zebrafish embryos were anesthetized by incubation in E2 embryo medium containing 40 mg/L tricaine (Sigma-Aldrich, E10521) until they no longer responded to tail stimulation. Embryos were euthanized at the completion of experiments by placement in a 500 mg/L solution of tricaine for at least 30 minutes. Male C57BL/6J mice aged 3-12 weeks were maintained under a 12-hour light/dark cycle at a constant temperature of 22°C. For sample collection, inhalational anesthesia with 4% isoflurane (Sigma-Aldrich, 1349003) was administered via a precision vaporizer in an induction chamber, followed by the use of a nose cone. Mice were then euthanized by exposure to 100% carbon dioxide at 5 PSI for a minimum of 3 minutes, at a displacement rate of 30% chamber volume per minute, in a cage or euthanasia chamber. Animals were left undisturbed after cessation of respiration for an additional 15 minutes.

### Plasmid constructs

To express RFP-F, the open reading frame (ORF) of monomeric RFP (mRFP) from pmRFP-C1 ^75^ was fused to the c-Ha-Ras farnesylation signal ^34, 35^ via PCR. For GFP-UtrCH expression, the first 783 base pairs of the human utrophin coding sequence ^26, 41^ were PCR-amplified and inserted in-frame into the BspEI-XhoI sites of pEGFP-C1. For *in vitro* mRNA transcription, the ORFs for H2B-GFP ^47^, H2B-RFP ^75^, RFP-F, GFP-hUtrCH, and EMTB-GFP ^46^ were subcloned into pCS2^+^. For heat shock-inducible expression of RFP or RFP-p50, the ORFs were PCR-amplified and inserted into the BamHI-NheI sites of pT2KhspGFF ^50^. pT2KhspGFF and pCS-TP, which encodes a transposase required for stable integration of pT2KhspGFF into genomic DNA ^76^, were generously provided by Dr. K. Kawakami (Graduate University for Advanced Studies, Japan). All plasmids containing PCR-amplified fragments were confirmed by sequencing.

### Zebrafish microinjection

*In vitro* mRNA synthesis was performed using the mMessage mMachine SP6 kit (Thermo Fisher, AM1340). For yolk injections, 5 nL of transcribed mRNA was injected, while for cytoplasmic injections, 1 nL of a plasmid-mRNA mixture was delivered into fertilized eggs. Phenol red (0.02%) was included as a tracer. For yolk injections, mRNAs encoding H2B-GFP, H2B-RFP, GFP-hUtrCH, or EMTB-GFP were used at a concentration of 50 ng/µL, while RFP-F mRNA was injected at 75 ng/µL. For cytoplasmic injections, a mixture containing H2B-GFP mRNA (20 ng/µL), TP mRNA (30 ng/µL), and a heat shock-inducible plasmid (30 ng/µL) was used. To induce gene expression, heat shock was applied for one hour at 39°C in a water bath starting at approximately 20 hpf.

### BrdU incorporation

For mouse experiments, C57BL/6 mice received intraperitoneal injections of BrdU at a dose of 100 µg per gram of body weight, followed by sacrifice at different time points. The epididymides were dissected and fixed overnight in 4% paraformaldehyde (PFA) in PBS at 4°C. BrdU labeling in zebrafish was performed as previously described^77^. Briefly, at 24 hpf, embryos were dechorionated and incubated in 10 mM BrdU (Millipore Sigma, B5002) in embryo medium for 20 minutes on ice. They were then transferred to warm medium and incubated for an additional 40 to 140 minutes at 28.5°C. After incubation, embryos were fixed overnight in PFA and subsequently dehydrated in graded methanol in PBT (PBS containing 0.1% Tween-20).

### Immunofluorescence microscopy

Mouse epididymal cryosections were prepared at a thickness of 10 µm and processed for staining as previously described ^78^. Briefly, tissue samples were blocked in PBS containing 10% goat serum and 0.5% Triton X-100 for one hour, followed by overnight incubation with primary antibodies at 4°C. After extensive washing, secondary antibodies were applied for 1.5 hours at room temperature. For BrdU staining, cryosections were denatured in 2 M HCl for 15 minutes at 37°C, neutralized in 0.1 M Na_2_B_4_O_7_ for 10 minutes at room temperature, and rinsed three times in PBS before incubation with an anti-BrdU antibody at a 1:1000 dilution. Whole-mount immunostaining of zebrafish embryos followed established protocols ^79^. BrdU staining was performed by digesting rehydrated embryos with proteinase K (10 µg/mL, Fisher Scientific, BP1700-100) for 10 minutes at room temperature, followed by fixation in PFA for 20 minutes. Embryos were incubated in 2 M HCl for one hour, washed three times in PBT, and then subjected to immunostaining ^77^. The primary antibodies used in this study included mouse monoclonal antibodies against α-tubulin (Millipore Sigma, T6199) and BrdU (Millipore Sigma, SAB4700630), rabbit antibodies against phospho-histone H3 (Ser10) (Cell Signaling Technology, 9701), phospho-E-cadherin (Ser838/840; Novus Biologicals, NBP3-12922), and β-catenin (Millipore Sigma, C2206), as well as a rat monoclonal antibody against E-cadherin (Abcam, ab11512). Secondary antibodies (ThermoFisher) used in this study were Alexa Fluor 488 goat anti-rabbit (A11034), goat anti-rat (A11006) and goat anti-mouse (A11029), and Alexa Fluor 546 goat anti-rabbit (A11035) and goat anti-mouse (A11030). Phalloidin-TRITC (P1951) and DAPI (D9542) were from Millipore Sigma. Fixed samples were imaged using Leica TCS SP5 or Zeiss LSM 800 confocal microscope.

### Live cell imaging

Embryos were first screened at 22 hpf under a Zeiss Axio Zoom V16 fluorescent dissecting microscope to identify those with GFP or RFP expression in the otic vesicles. Selected embryos were dechorionated, anesthetized with tricaine, mounted in 1% low-melting point agarose (Fisher Scientific, BP1360-100) within a glass-bottom Petri dish (MatTek Life Sciences, P35G-1.5-14-C), and covered with E2 embryo medium containing tricaine and 0.003% phenylthiourea (Millipore Sigma, P7629). For experiments involving drug treatments, blebbistatin (100 µM, Millipore Sigma, B0560), nocodazole (0.5 µg/ml, Millipore Sigma, M1404), or DMSO (Millipore Sigma, D8418) as the solvent control was added to both the medium and agarose. Embryos were positioned so that one otic vesicle was in close proximity to the glass bottom. Time-lapse imaging was conducted at 2-minute intervals over a period of 216 to 430 minutes using a 20× NA 0.7 objective on a Leica TCS SP5 or Zeiss LSM 800 inverted confocal microscope equipped with a heating hood.

### Statistical analysis of nuclear migration

Nuclear displacement was quantified as the distance between the bottom edge of the nucleus and the basal membrane^26^. Measurements were taken at 2-minute intervals using ImageJ software. Prometaphase was identified by condensed but dispersed chromosomes accompanied by strong pH3 signal. Metaphase was defined by condensed chromosomes aligned at the cell center with strong pH3 signal. Anaphase was defined by separated chromosomes with reduced pH3 signal, whereas telophase nuclei showed no detectable pH3 signal. Cell rounding was defined as a mitotic shape change in which the main cell body becomes spherical while retaining a thin basal projection attached to the basal lamina, thereby separating the bulk of the cell body from the basal membrane. In contrast, a diamond-shaped morphology was defined by cells that maintained a broad basal attachment to the basal lamina. Quantification values are reported as mean ± S.D. Two-tailed Mann-Whitney U test was applied for small sample sizes (3-5 replicates). Nested t test was applied when involving hierarchical data (cells nested within animals).

## Data availability

Data generated in this study are included in the Supplementary file.

## Acknowledgements

We thank A. Afolalu, C. Shapiro, S. Hosten, and C. Quaies for fish care; Dr. Koichi Kawakami (National Institute of Genetics and Department of Genetics, Graduate University for Advanced Studies, Japan) for providing the pT2KhspGFF and pCS-TP constructs; Yanan Xu, Zhili Wu, and Yiping Li for technical support; and Xueliang Zhu for guidance and comments on the manuscript.

## Author contributions

Conceptualization: J.C.

Methodology: Y.X. and J.C.

Investigation: Y.X., B.P., A.G.C.Y., M.M., and M.Q.

Resources: G.S.P. and J.C.

Writing and editing: Y.X. and J.C.

Funding Acquisition: J.C.

## Supplementary Information

### Legends for supplementary figures

**Supplementary Fig. S1. Nocodazole treatment does not affect G2 entry.** Zebrafish otic vesicle was stained for α-tubulin (green), pH3 (magenta), and DAPI (blue) after 2, 4, and 8 h treatment of DMSO or Nocodazole. Scale bar, 20 µm

### Legends for supplementary videos

**Supplementary Video S1. Nuclear migration and mitosis in zebrafish otic vesicle.** Fertilized embryos were microinjected with *in vitro* transcribed mRNA to express H2B-GFP, labeling chromatin. Imaging of the otic vesicle began at approximately 22 hpf and continued for 170 minutes at 2-minute intervals.

**Supplementary Video S2. Morphological changes of a mitotic cell in the zebrafish otic vesicle.** H2B-GFP (green) labels chromatin, while RFP-F (red or gray) labels the plasma membrane. The arrow highlights the nucleus, and arrowheads indicate the lateral plasma membrane.

**Supplementary Video S3. F-actin dynamics in a mitotic cell of the zebrafish otic vesicle.** H2B-RFP (red) labels chromatin, and GFP-hUtCH (green or gray) labels F-actin. The arrow marks the nucleus, while arrowheads point to the lateral plasma membrane.

**Supplementary Video S4. Nuclear migration and mitosis in the zebrafish otic vesicle following DMSO treatment.** Embryos were microinjected with mRNA to express H2B-RFP (red) and GFP-hUtrCH (green). Imaging started approximately 20 minutes after DMSO was added at 22 hpf.

**Supplementary Video S5. Nuclear migration and mitosis in the zebrafish otic vesicle following Blebbistatin treatment.** Embryos were microinjected with mRNA to express H2B-RFP (red) and GFP-hUtrCH (green). Imaging began approximately 25 minutes after Blebbistatin was added at 22 hpf.

**Supplementary Video S6. Microtubule dynamics in an otic vesicle cell entering the cell cycle.** H2B-RFP (red) labels chromatin, and EMTB-GFP (green) labels stable microtubules.

**Supplementary Video S7. Control otic vesicle treated with DMSO.** Embryos were microinjected with mRNA to express H2B-GFP (green) and RFP-F (red). Imaging commenced approximately 30 minutes after DMSO was added at 22 hpf.

**Supplementary Video S8. Otic vesicle morphology following nocodazole treatment.** Embryos were microinjected with mRNA to express H2B-GFP (green) and RFP-F (red). Imaging started approximately 30 minutes after nocodazole was added at 22 hpf.

**Supplementary Video S9. A control otic vesicle overexpressing RFP**. The zebrafish embryo was microinjected with both mRNA to express H2B-GFP (green) and a plasmid to overexpress RFP (red) in a heat shock-inducible manner. The heat shock treatment was performed at 20 hpf for 1hr. The otic vesicle was imaged at 1.5 hr after the heat shock was done.

**Supplementary Video S10. A representative otic vesicle overexpressing RFP-p50**. The zebrafish embryo was microinjected with both mRNA to express H2B-GFP (green) and a plasmid to overexpress RFP-p50 (red) in a heat shock-inducible manner. The heat shock treatment was performed at 20 hpf for 1hr. The otic vesicle was imaged at 1.5 hr after the heat shock was done.

**Supplementary file:** Quantification results used to generate plots.

