## Supplementary figures and images for "Zebrafish otic vesicle and mouse epididymis as model systems for studying columnar epithelial cell division"

### Figure S1

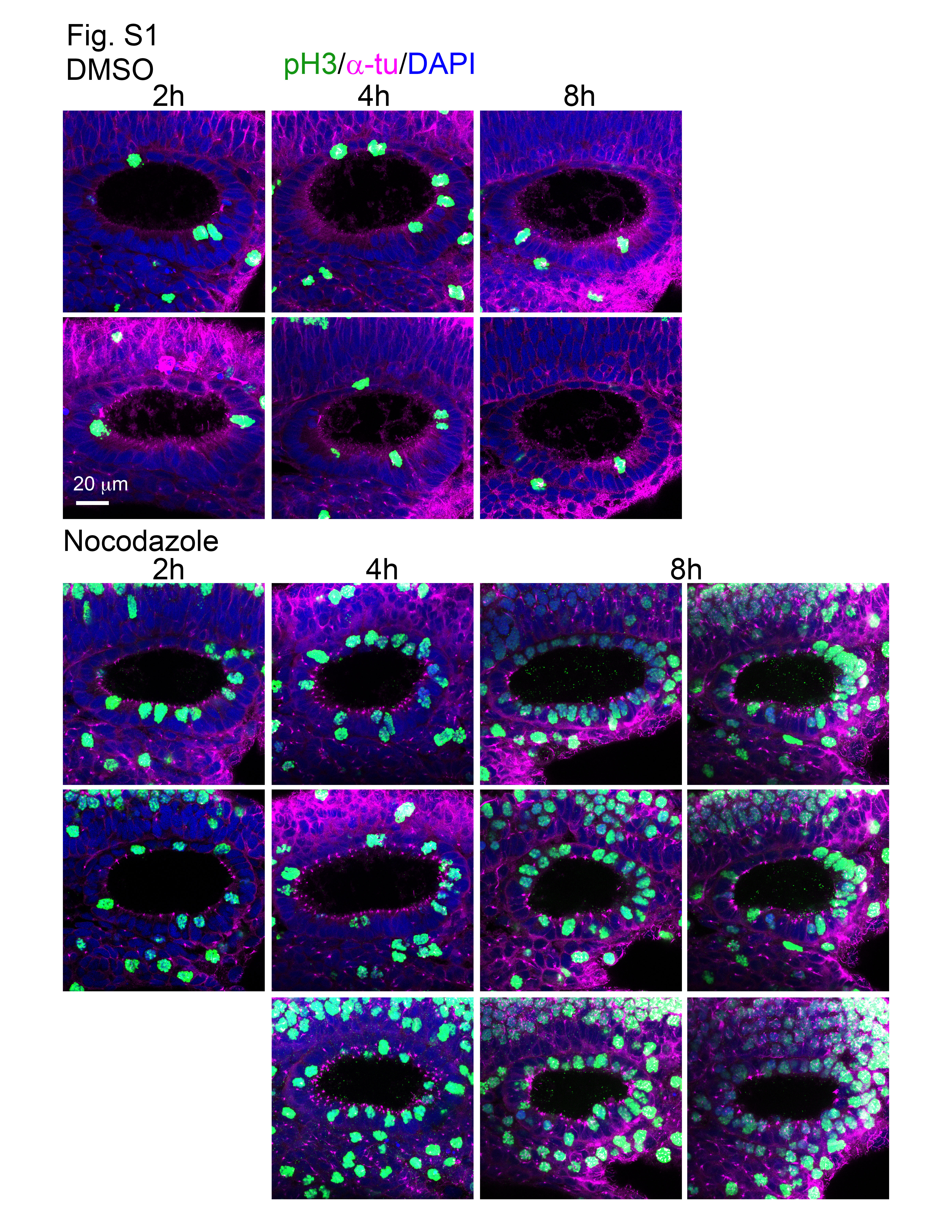
